# *VPS13D* mutations affect mitochondrial homeostasis and locomotion in *Caenorhabditis elegans*

**DOI:** 10.1101/2025.01.22.634397

**Authors:** Xiaomeng Yin, Ruoxi Wang, Andrea Thackeray, Eric H. Baehrecke, Mark J. Alkema

**Author notes:** Correspondence to Eric Baehrecke, and Mark Alkema.

## Abstract

Mitochondria control cellular metabolism, serve as hubs for signaling and organelle communication, and are important for the health and survival of cells. *VPS13D* encodes a cytoplasmic lipid transfer protein that regulates mitochondrial morphology, mitochondria and endoplasmic reticulum (ER) contact, quality control of mitochondria. *VPS13D* mutations have been reported in patients displaying ataxic and spastic gait disorders with variable age of onset. Here we used CRISPR/Cas9 gene editing to create *VPS13D* related-spinocerebellar ataxia-4 (SCAR4) missense mutations and C-terminal deletion in *VPS13D*’s orthologue *vps-13D* in *C. elegans*. Consistent with SCAR4 patient movement disorders and mitochondrial dysfunction, *vps-13D* mutant worms exhibit locomotion defects and abnormal mitochondrial morphology. Importantly, animals with a *vps-13D* deletion or a N3017I missense mutation exhibited an increase in mitochondrial unfolded protein response (UPR^mt^). The cellular and behavioral changes caused by *VPS13D* mutations in *C. elegans* advance the development of animal models that are needed to study SCAR4 pathogenesis.

## Introduction

Mitochondria are dynamic organelles that play an important role in cellular bioenergetics and viability. The mitochondrial network undergoes continuous fusion and fission to maintain its health. In addition, mitochondria form dynamic contacts with other organelles, especially the endoplasmic reticulum, to participate in various biological processes. Damaged mitochondria are selectively cleared by autophagy, and alterations in mitochondrial removal by autophagy have been associated with neurological disorders, including Alzheimer’s disease, Parkinson’s disease, Huntington’s disease, and amyotrophic lateral sclerosis (Collier et al. 2023).

Vacuolar protein sorting-associated protein 13D, encoded by the *VPS13D* gene, is a key regulator of mitochondrial processes, including mitochondrial clearance, mitochondrial morphology, and inter-organelle contacts (Anding et al. 2018; Shen et al. 2021b; Guillen-Samander et al. 2021). VPS13D is a member of the conserved VPS13 protein family, which facilitates membrane contacts and transport lipids between organelles (Leonzino et al. 2021). Unlike the other three human VPS13 proteins (VPS13A–C), VPS13D possesses a unique ubiquitin-associated (UBA) domain that has been shown to interact with K63 ubiquitin chains and participate in mitochondrial health (Anding et al. 2018; Leonzino et al. 2021). Mutations in the *VPS13A–C* genes have been linked to specific neurological disorders, including chorea-acanthocytosis, Cohen syndrome, and early-onset Parkinson’s disease. Importantly, biallelic pathogenic variants in the *VPS13D* gene have also been associated with spastic ataxia and spastic paraplegia (Seong et al. 2018; Gauthier et al. 2018).

VPS13D movement disorder, also referred to as autosomal recessive spinocerebellar ataxia-4 (SCAR4; (Swartz et al. 2002; Seong et al. 2018)), manifests as a hyperkinetic movement disorder characterized by dystonia, chorea, and/or ataxia. This disorder is often accompanied with varying degrees of developmental delay and cognitive impairment (Meijer 2019). In the majority of the reported 42 cases, individuals have a combination of a loss-of-function variant and a seemingly milder variant, which can either be a missense mutation or a milder splicing variant (Seong et al. 2018; Gauthier et al. 2018; Koh et al. 2020; Petry-Schmelzer et al. 2021; Wang Yu 2021; Huang and Fan 2022; Oztop-Cakmak et al. 2022; Durand et al. 2022; Pauly et al. 2023; Baker et al. 2023; Harada et al. 2024; Sultan et al. 2024; Nordli and Galan 2023; Kistol et al. 2023; Kistol et al. 2024; Algahtani et al. 2024; Lee et al. 2020). Both *VPS13D*-deficient human and *Drosophila* cells exhibit notable abnormalities in mitochondrial morphology and clearance (Seong et al. 2018; Anding et al. 2018). In addition, a genetic screen for altered mitochondria in *C. elegans* identified *C25H3.11* (*vps-13D*) (Rolland and Conradt 2022). Recent studies have linked *VPS13D* to Leigh syndrome (Kistol et al. 2023; Lee et al. 2020) and the Parkinson’s disease gene *PINK1* (Shen et al. 2021a) suggesting that this gene has relevance to a broader spectrum of neurological diseases.

Strong loss-of-function *VPS13D* alleles are lethal in *Drosophila* (Anding et al. 2018) and mice (Skarnes et al. 2011), thus presenting a challenge to model this disease in an intact animal. In this study, we used CRISPR/Cas9 gene editing techniques to create SCAR4 patient-specific mutations in the *C. elegans* orthologue of the human *VPS13D* gene: *vps-13D*. We analyzed the effects of these SCAR4 patient-specific mutations in *vps-13D* on worm fecundity, behavior, and mitochondrial morphology. We find that *vps-13D* deletion mutants display maternal effect sterility and locomotion defects. *vps-13D* deletions and the N3017I missense mutation displayed disrupted mitochondrial morphology and resulted in different degrees of activated mitochondrial unfolded protein response (UPR^mt^). Furthermore, we identified a functional link between *vps-13D* and the mitochondria fusion regulator *fzo-1* in the same pathway to regulate mitochondrial dynamics and homeostasis. The combined biological changes caused by *VPS13D* mutations in *C. elegans* advance the development of a new animal model to study SCAR4 pathogenesis.

## Materials and methods

### C. elegans strains and GFP fusion construction

All *C. elegans* strains were maintained at 22°C on nematode growth medium (NGM) plates seeded with *Escherichia coli* OP50 bacteria. The wild-type strain was Bristol N2. Sterile mutant *vps-13D* strains were balanced with the *mIn1* balancer chromosome (Edgley and Riddle 2001). The *vps-13D* mutant strains were crossed with the following transgenes: *zcIs14* [*Pmyo-3::GFP* (*mit*)] (Benedetti et al. 2006), *zcIs13* [*Phsp-6::GFP + lin-15* (*+*)] (Yoneda et al. 2004; Yang et al. 2022). A full list of strains used in this study is shown in Supplementary Table S1. The *Pvps-13D*::*GFP* transcriptional reporter was generated using the following primers: F(5’-ACAGGATCCTCGCATACAATCACATCGTC-3’) and R(5’-ACAGGTACCTGTCCAGGAATTGTGGTATC-3’). These primers were used to amplify a 3.5 kb fragment corresponding to the upstream promoter sequence (including exon 1 and part of exon 2) of *vps-13D*. The PCR product was digested by BamHI and KpnI restriction enzymes and inserted into pPD95.75 vector. The *Pvps-13D*::*GFP* construct was microinjected at 50 ng/µl into temperature-sensitive *lin-15*(*n765ts*) mutant animals along with the *lin-15* (*+*) rescuing plasmid (pL15EK) at 80 ng/µl and pBSK DNA at 80 ng/µl. Transgenics were selected at 22°C based on a non-Muv (Multivulva) phenotype and GFP fluorescence.

### CRISPR/Cas9 design and gene editing

The Bristol N2 strain was used as the wild-type strain for all CRISPR/Cas9 editing of *vps-13D*. Site-specific crRNAs and a repair template donor ssODNs were manually selected and generated by IDT (Integrated DNA Technologies, Inc.) who also provided the tracrRNA. Injection mixtures were prepared following established protocols (Ghanta et al. 2021) and *rol-6* was used as a co-CRISPR selection marker. All crRNAs, ssODNs, and PCR screening sequences are reported in Supplementary Table S2-S4. Correct substitution or deletion sequences were confirmed via Sanger sequencing. Sequencing of C-terminal deletion *ΔC* (*zf197*) revealed a 1218-bp deletion (LGII 5670774–5671991) with a short 20-bp insertion positioned between the two breakpoints. The CRISPR-designed strains underwent four rounds of outcrossing to the N2 strain.

### Brood size assay

Ten L4 animals of each genotype were picked and separated onto individual plates. Animals were transferred onto new plates every day until the cessation of egg-laying. Plates were counted for the total number of eggs after removal of the parent animal.

### Larval development analysis

Five adult animals from each genotype were transferred to a new plate. After 2-hours, adult animals were removed, and the eggs were left to develop for over 96 hours. Two *vps-13D* deletion mutant strains were maintained as balanced heterozygotes using the *mIn1* [*dpy-10* (*e128*) *mIs14*] balancer. This balancer includes an integrated pharyngeal GFP reporter expressed in a semi-dominant manner. Consequently, the offspring segregate into wild type with a dim GFP signal, Dpy with a bright GFP signal (*mIn1* homozygotes), and non-GFP *vps-13D* deletion homozygotes. For the *vps-13D* deletion strains only non-GFP homozygotes were evaluated at each developmental stage. The developmental timing was calculated as the proportion of larvae among the total number of hatched embryos that reached each specific developmental stage. The developmental stage of each individual was determined based on body size and stage-specific morphological features.

### Multi-Worm Tracker assay

10 synchronized 1-day old or 3-day old (24 hours or three days post the L4 stage) worms were placed on a thin lawn *E. coli* OP50 on a medium (5 cm) NGM plate. The plate was then placed in the Multi-Worm Tracker (MWT) (https://sourceforge.net/projects/mwt/, (Swierczek et al. 2011)) and recorded for 10 minutes. The Multi-Worm Tracker package includes real-time image analysis software and behavioral parameter measurement software, Choreography. Tracking and analysis were conducted following previous studies (Huang et al. 2019; Florman and Alkema 2022; Kang et al. 2024). The average speed of the population was measured over 5 minutes after a 5-minute acclimation period. Experiments were analyzed using custom MATLAB (MathWorks, Inc.) scripts to interface with Choreography analysis program. The MATLAB scripts used in this study are available at https://github.com/jeremyflorman/Tracker_GUI. Analysis was limited to objects that had been tracked for a minimum of 20 seconds and had moved a minimum of 5 body lengths. For the analysis of single worm tracks, a plate containing a single 3-day old animal was recorded for 10 minutes with images captured every 2 seconds. The sequential images from the final 5 minutes post-acclimation were stitched and processed in ImageJ to generate a continuous worm movie track.

### Thrashing assay

10 age-synchronized young adult animals were transferred to a fresh unseeded plate to remove residual bacteria. Individual animals were subsequently transferred to M9 buffer in a tiny plate. After 15 seconds acclimatization, the number of completed thrashes during 1 minute were counted per animal using a hand counter.

### Gene expression knockdown via RNAi treatment

*fzo-1* RNAi bacteria clones, obtained from the Ahringer library (Kamath and Ahringer 2003), were selected by ampicillin (100 mg/ml) and tetracycline (12.5 mg/ml) and verified by DNA sequencing. The control L4440 or *fzo-1* RNAi bacteria grown at 37°C overnight in LB with ampicillin (100 mg/ml), were concentrated (4X) and seeded on RNAi NGM plates that contain 6 mM IPTG and 100 mg/ml ampicillin. Young adult P0 animals were placed on the RNAi plates and allowed to produce offspring. L4 larvae of F1 progeny were transferred to a new RNAi plate. After two generations of RNAi-treatment, L4 larvae of F2 animals were analyzed.

### Imaging and fluorescence quantification

To assess the sterile phenotype, adult wild-type animals and *vps-13D* deletion mutant animals were placed on 2% agarose pads containing 60 mM sodium azide. Adult sterile animals were identified by the absence of embryos or oocytes in the uterus through DIC imaging using a Zeiss LSM 700 microscope. To explore the expression pattern of *vps-13D*, adult animals expressing a *Pvps-13D*::*GFP* reporter were mounted on 2% agarose pads containing 60 mM sodium azide. Images were captured using a Zeiss LSM 700 confocal microscope. To test the impact of *vps-13D* mutations on mitochondrial morphology, L4 animals expressing *zcIs14* transgene were mounted on 2% agarose pads containing 10 mM levamisole. Images were recorded using a Zeiss LSM 700 confocal microscope. For each genotype mitochondrial morphology in body wall muscles in the middle of the worm body was scored for 10∼15 animals in total. To test the effects of *vps-13D* mutations on *hsp-6* expression of *C. elegans*, six L4 animals expressing *Phsp-6*::*GFP* grown on *E. coli* OP50 were mounted together on 2% agarose pads containing 10 mM levamisole and 18∼36 animals in total. The GFP images were acquired with an Axioimager Z1 microscope (Zeiss) at 10x magnification, and the maximum fluorescence intensity was quantified using ImageJ FIJI software.

### Mitochondrial morphology analysis

Mitochondrial images were segmented and quantified using the Mitochondrial Segmentation Network (MitoSegNet) toolbox, a deep learning-based tool that includes the MitoS segmentation tool and the MitoA analysis tool (Fischer et al. 2020). Documentation for the MitoS and MitoA tools are available at https://github.com/mitosegnet, and the MitoSegNet segmentation model, along with the MitoA analysis and MitoS segmentation tools (GPU/CPU versions) for Linux and Windows, can be accessed at https://zenodo.org/search?page=1&size=20&q=mitosegnet. Initially, 8-bit raw images were segmented using the pretrained mitochondria-specific “basic mode” (MitoSegNet model) in the MitoS tool (CPU version). Upon completion, the program generated a prediction folder containing the segmented images. The segmented images and corresponding raw images were then subjected to quantitative measurement using the MitoA tool, which evaluates morphological features for each object and generates summary statistics for all object features in each image. The major axis lengths were determined by measuring the lengths of the line segment connecting the two vertices of an ellipse fitted around an object in the MitoA tool.

### Statistical analysis

All statistical analysis and graph construction were performed using GraphPad Prism 8 software. Results are presented as standard error of the mean (SEM) from at least three independent experiments. Statistical comparisons were performed using ANOVA or Kruskal-Wallis H test with Dunnett’s correction for multiple samples. Significance was determined when P-value < 0.05.

## Results

### C. elegans vps-13D encodes an ortholog of VPS13D

To explore the evolutionary relationships of the VPS13D proteins among various species, we first conducted a phylogenetic analysis. VPS13D proteins from different species formed a well-supported clade, indicating that they evolved from a common ancestor (Fig. 1A). The *C. elegans C25H3.11* gene encodes a protein that shares the most significant similarity with VPS13D. The predicted Ricin B-type lectin domain-containing protein shares 36% similarity and 22% identity with the human VPS13D protein (Table S5). Furthermore, the highest sequence similarity between the two proteins was observed in the C-terminal region, which contains the VPS13 adaptor binding (VAB) domain (Bean et al. 2018) (also known as SHR binding domain or WD40-like region (Kumar et al. 2018)) and a Dbl homology (DH)-like domain (InterPro) (Fig. 1B). Given that *C25H3.11* gene is the closest *C. elegans* orthologue of the human *VPS13D* gene, we hereafter refer to *C25H3.11* as the *C. elegans vps-13D* gene. In humans, 14 of 30 reported families carry at least one missense mutation in the C-terminal region following the ubiquitin-associated (UBA) domain, and 5 of 8 reported homozygous missense mutations in patients are also located in this region (Seong et al. 2018; Gauthier et al. 2018; Koh et al. 2020; Petry-Schmelzer et al. 2021; Wang Yu 2021; Huang and Fan 2022; Oztop-Cakmak et al. 2022; Durand et al. 2022; Pauly et al. 2023; Baker et al. 2023; Harada et al. 2024; Sultan et al. 2024; Nordli and Galan 2023; Kistol et al. 2023; Kistol et al. 2024; Algahtani et al. 2024; Lee et al. 2020). Several of these mutation sites associated with human pathogenicity are highly conserved in the C-terminal region in *C. elegans* VPS13D protein (Fig. 1C).

**Fig. 1.**
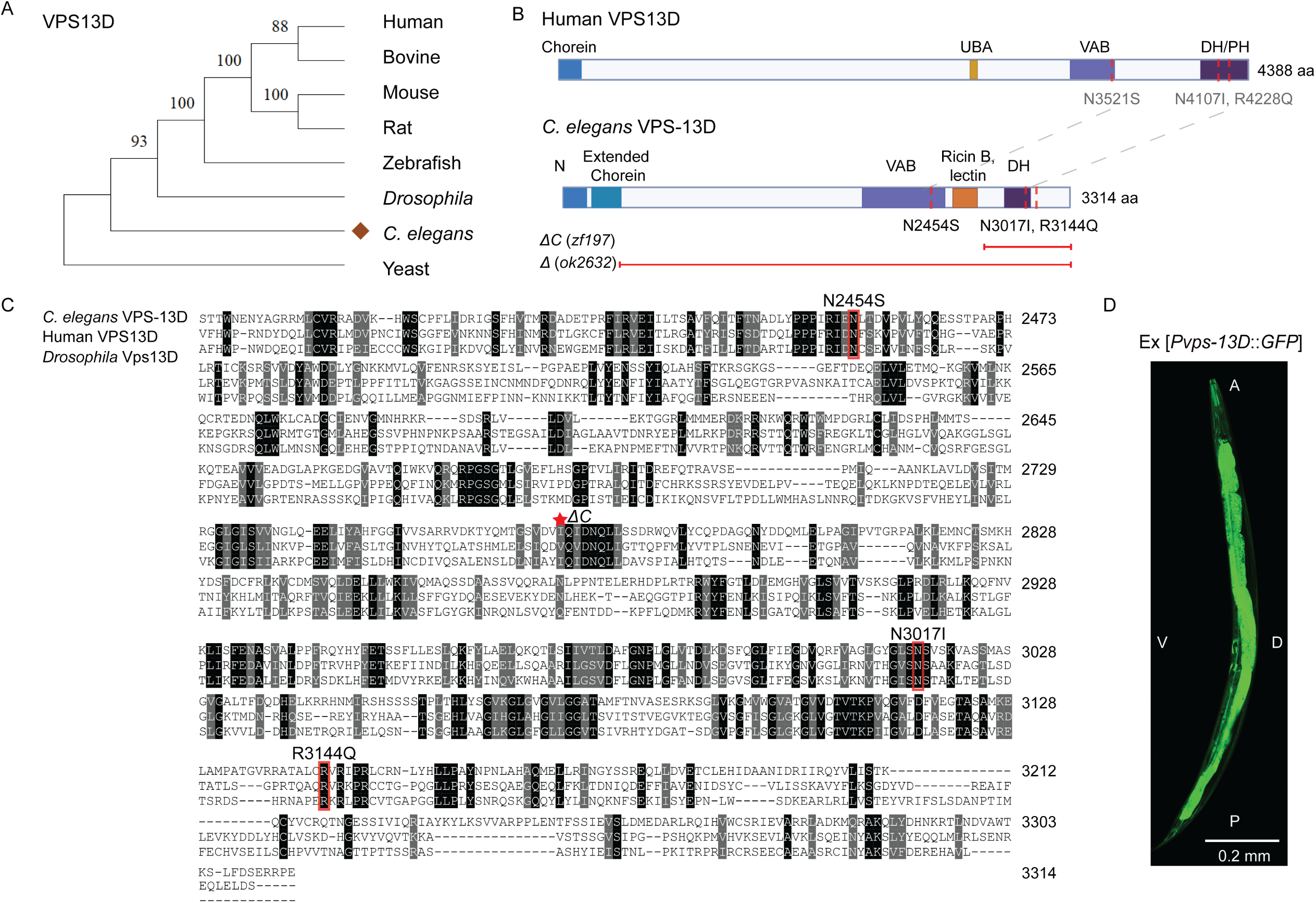
*VPS13D* orthologue *vps-13D* is conserved in *C. elegans* (A) Phylogenetic tree of VPS13D proteins. VPS13D protein sequences from various species were retrieved from UniProt. The evolutionary history was inferred using the neighbor-joining method. The percentage of replicate trees in which the associated taxa clustered together in the bootstrap test (1000 replicates) are shown next to the branches. Evolutionary analyses were conducted in MEGA11. The species depicted from top to bottom are Human (UniProt identifier: Q5THJ4), Bovine (UniProt identifier: E1BIF6), Mouse (UniProt identifier: B1ART2), Rat (UniProt identifier: A0A8I6G572), Zebrafish (UniProt identifier: A0A8M1QUR7), *Drosophila* (UniProt identifier: Q9VU08), *C. elegans* (labeled as brown diamond, UniProt identifier: A0A2K5ATR5) and Yeast (UniProt identifier: Q07878). (B) Human VPS13D protein topology (modified from (Guillen-Samander et al. 2021) and predicted domains of the *C. elegans* VPS-13D protein with disease-related missense mutations labeled in dotted vertical lines and deletions annotated with horizontal lines. Domains and abbreviations are as follows: N, Chorein N-terminal domain; Chorein domain; Extended Chorein domain; UBA, Ubiquitin-associated domain; VAB, Vps13 Adaptor Binding/SHR-Binding/WD40-like domain; Ricin B-type lectin domain; DH, Dbl homology-like domain; and PH, Pleckstrin homology domain. (C) Sequence alignment of C terminal region between *C. elegans* VPS-13D, human VPS13D, and fly Vps13D protein by Clustal Omega. The three missense mutations enrolled in this study are highlighted with the red box, and the start residue of *vps-13D*(*ΔC*) deletion is labeled with the red asterisk. (D) Representative fluorescence image of adult hermaphrodites expressing *Pvps-13D*::*GFP* under the 3570bp promoter (including exon 1 and part of exon 2). A, anterior; P, posterior; V, ventral side; D, dorsal side. The scale bar is 0.2 mm.

We next focused on three conserved residues within the C-terminal region of VPS13D protein that are reported to be mutated in early onset SCAR4 patients. We used CRISPR/Cas9 gene editing to create these variants in *vps-13D* (*N2454S*(*zf194*), *N3017I*(*zf195*) and *R3144Q*(*zf196*)) (Table 1). In addition, we generated a C-terminal deletion *ΔC*(*zf197*) that encompassed 1218 base pairs, resulting in a truncation of the VPS13D protein (Fig. 1B). We also obtained a *vps-13D* deletion allele, *Δ*(*ok2632*), which contains a 1740 bp deletion removing part of exons 3 and 4, resulting in a premature stop and thus most likely represents a null allele. To analyze the expression of *vps-13D*, we investigated its expression pattern by generating a transcriptional fusion of GFP reporter with 3.5kb *vps-13D* upstream promoter fragment (including exon 1 and part of exon 2).

**Table 1.** *VPS13D* pathogenic missense mutations in this study.

| *C. elegans*<br>mutations<br>(*vps-13D* a.1) | *VPS13D* pathogenic<br>mutations<br>(NM_015378) | Clinical diagnosis<br>of patients | Onset age |
| --- | --- | --- | --- |
| N2454S<br>(*zf194*) | p.N3521S<br>(c.10562A>G) | Spastic ataxia with<br>mild intellectual<br>disability (2 patients) | 2y/4-5y |
| N3017I<br>(*zf195*) | p.N4107I<br>(c. 12320A>T) | Ataxia, neuropathy,<br>developmental delay,<br>seizures, and optic<br>atrophy | 1y |
| R3144Q<br>(*zf196*) | p.R4228Q<br>(c.12683G>A) | Ataxia, muscle<br>weakness, dystonia,<br>mild intellectual<br>disability | 1-2y |

Fluorescence from the *Pvps-13D*::*GFP* reporter was observed throughout the adult hermaphrodite, including intestine, hypodermis, muscle, glia and neurons (Fig. 1D). Our data indicate that *vps-13D* is expressed in most cell types consistently with RNA expression data sets (Taylor et al. 2021).

### *vps-13D* mutants exhibit impaired locomotion

Animals carrying either homozygous *vps-13D* deletion alleles (*Δ*(*ok2632*) or *ΔC*(*zf197*)) are viable. However, both homozygous *vps-13D*(*Δ*) and *vps-13D*(*ΔC*) animals display maternal effect sterility (Fig. 2A). Wild-type hermaphrodites produce oocytes that are fertilized in the spermatheca, resulting in the accumulation of developing embryos in the uterus. The germlines of either homozygous *vps-13D*(*Δ*) or *vps-13D*(*ΔC*) animals did not display visible oocytes or embryos (Fig. 2B). Due to their sterility, these two deletion strains were maintained as balanced heterozygotes but analyzed as homozygotes by selecting homozygous animals from a mixed population. In addition to fertility deficiencies, these two deletion mutants also exhibited a slower rate of development, taking 24∼48 hours longer than the wild type to reach the L4 stage (Fig. 2C). In contrast, three missense mutants *vps-13D*(*N2454S*), (*N3017I*), and (*R3144Q*) did not exhibit developmental arrest or delay and were able to produce a similar brood size compared to the wild type (Fig. 2A).

**Fig. 2.**
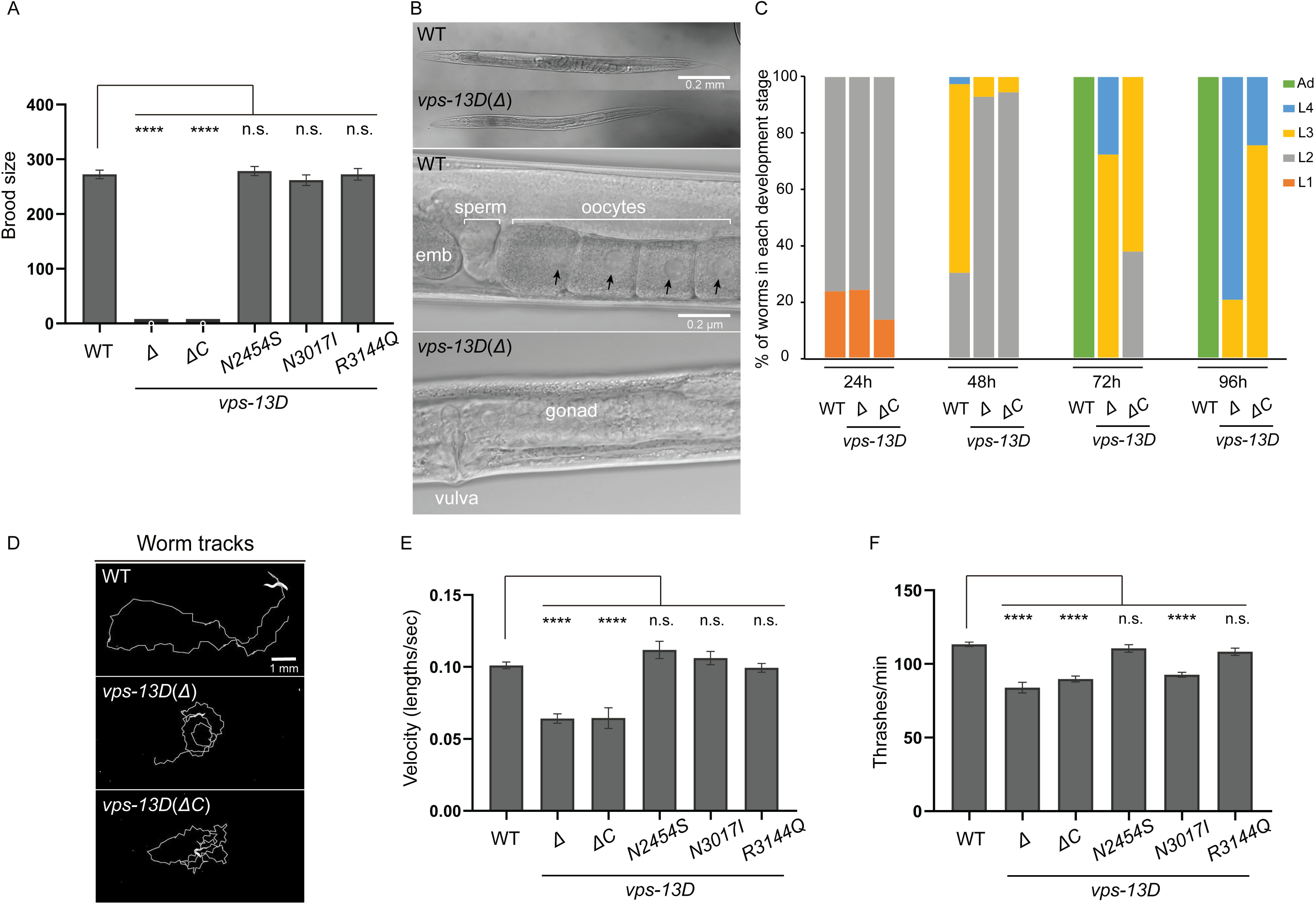
*vps-13D* deficiency affects worms’ fertility and locomotion. (A) Total brood size (n=30 broods) indicates the fertility ability of *C. elegans* to produce offspring in association with different *vps-13D* mutants. One-way ANOVA with Dunnett correction was used to compare the difference between wild-type and mutant animals. (B) DIC images of whole worm and gonad in hermaphrodites of the wild type and *vps-13D* (*Δ*) mutant strains. The scale bar in the whole worm image is 0.2 mm. The scale bar in the gonad image is 0.2 μm. Black arrows indicate oocyte nuclei. (C) The percentage of worms at the different developmental stages was determined for wild-type, *vps-13D*(*Δ*), and *vps-13D*(*ΔC*) mutant animals after 96 hours of growth from the embryonic stage at 22°C. (D) Individual animal tracks of 3-day-old wild type, *vps-13D*(*Δ*) and *vps-13D*(*ΔC*) mutant strains for the final 5 min recordings. The scale bar is 1 mm. (E) Comparison of average velocity for 3-day-old wild-type and *vps-13D* animals from the final 5 mins MWT recordings. The velocity has been normalized and calculated based on the worm body lengths of respective strains. The difference in average velocity between wild-type and mutants was analyzed by one-way ANOVA with Dunnett’s multiple comparison test. (F) Number of thrashes per minute in M9 liquid for 3-day-old wild type and *vps-13D* mutant animals (n=15-30). Differences between wild type and mutants were analyzed by One-way ANOVA with Dunnett correction for multiple comparisons.

Next, we examined whether *vps-13D* mutant strains possess motor defects. We conducted a population-level motility assay using an automated multi-worm tracking system (MWT) (Swierczek et al. 2011). We tracked wild type and *vps-13D* mutant strains for 5 minutes and extracted the behavioral dynamics of individual worms. One-day-old adult *vps-13D* mutants did not shown any difference compared to the wild-type (Fig. S1). However, *vps-13D* deletion animals displayed a noticeable movement impairment compared to wild-type animals. The average locomotion rate of 3-day-old wild-type animals was 0.101± 0.005 body lengths/sec, whereas either *vps-13D*(*Δ*) or *vps-13D*(*ΔC*) mutants displayed decreased locomotion rates of 0.064±0.010 and 0.065±0.012 body lengths/sec (Fig. 2D-E). The three *vps-13D* missense mutant strain locomotion rates were unaffected at day 3 of the adult stage (0.111±0.009, 0.106±0.010, 0.099±0.008 body lengths/sec, respectively, Fig. 2E). Furthermore, we analyzed thrashing rates of single worm in liquid M9 medium. The average thrashing rate of 3-day-old wild-type animals was 113.2±6.9 thrashes/min. Significantly, *vps-13D*(*Δ*) and *vps-13D*(*ΔC*) mutant strains showed a 20∼26% decrease in thrashing rates compared to control wild-type animals (83.8 ± 11.0 thrashes/min, 89.8±8.7 thrashes/min, respectively, Fig. 2F). Importantly, the thrashing rate of *vps-13D*(*N3017I*) mutant worms was 92.7±7.1 thrashes/min, showing an 18% reduction compared to control animals (Fig. 2F), where *vps-13D*(*N2454S*) and *vps-13D*(*R3144Q*) mutants exhibited similar thrashing rates to the wild type (110.5±11.0 thrashes/min, 108.3±9.5 thrashes/min, respectively). Combined, these results indicate that the *C. elegans vps-13D* mutations can result in locomotion defects.

### *vps-13D* mutants possess altered mitochondrial morphology

Dynamic changes in mitochondrial morphology are essential for mitochondria health and homeostasis. Mitochondrial fusion and fission not only affect mitochondria morphology but also regulate multiple mitochondria biological processes, including mitochondria quality control and clearance by mitophagy (Pickles et al. 2018). Previous studies indicate that VPS13D affects mitochondrial morphology and mitophagy in both human and *Drosophila* cells (Anding et al. 2018). To test if *vps-13D* mutants exhibit a similar function to maintain mitochondrial morphology, we used a transgenic line *zcIs14* ([*myo-3::GFP* (*mit*)]) that labels mitochondria in the body wall muscles (Benedetti et al. 2006) and quantified the mitochondrial morphology using MitoSegNet (Fischer et al. 2020). Although morphologies between different mutants were variable and possibly reflected allele strength, mutant strains of *vps-13D* displayed changes in mitochondrial morphology at the L4 larval stage compared to the wild type (Fig. 3A). Specifically, all homozygous *vps-13D* mutant worms possessed shorter mitochondria compared to control wild-type worms, as determined by the major axis length (Fig. 3A and 3B). In wild-type worms, the major axis length was 31.4±11.6 pixels. By contrast, *vps-13D*(*Δ*) and *vps-13D*(*ΔC*) mutant strains displayed significantly reduced major axis lengths of 18.1±3.7 pixels and 19.0±5.4 pixels (Fig. 3B). Similarly, the major axis length was also reduced in the three *vps-13D* missense mutant strains, measuring 19.7±5.7, 18.0±3.9, 22.3±9.5 pixels, respectively (Fig. 3B). These data are consistent with a recent report (Rolland and Conradt 2022). In addition, both *vps-13D*(*Δ*) and *vps-13D*(*ΔC*) mutant strains demonstrated a notable increase in mitochondrial area measuring 154.0±32.6 pixels and 144.7±30.2 pixels, respectively, compared to 113.5±19.6 pixels in wild-type worms.

**Fig. 3.**
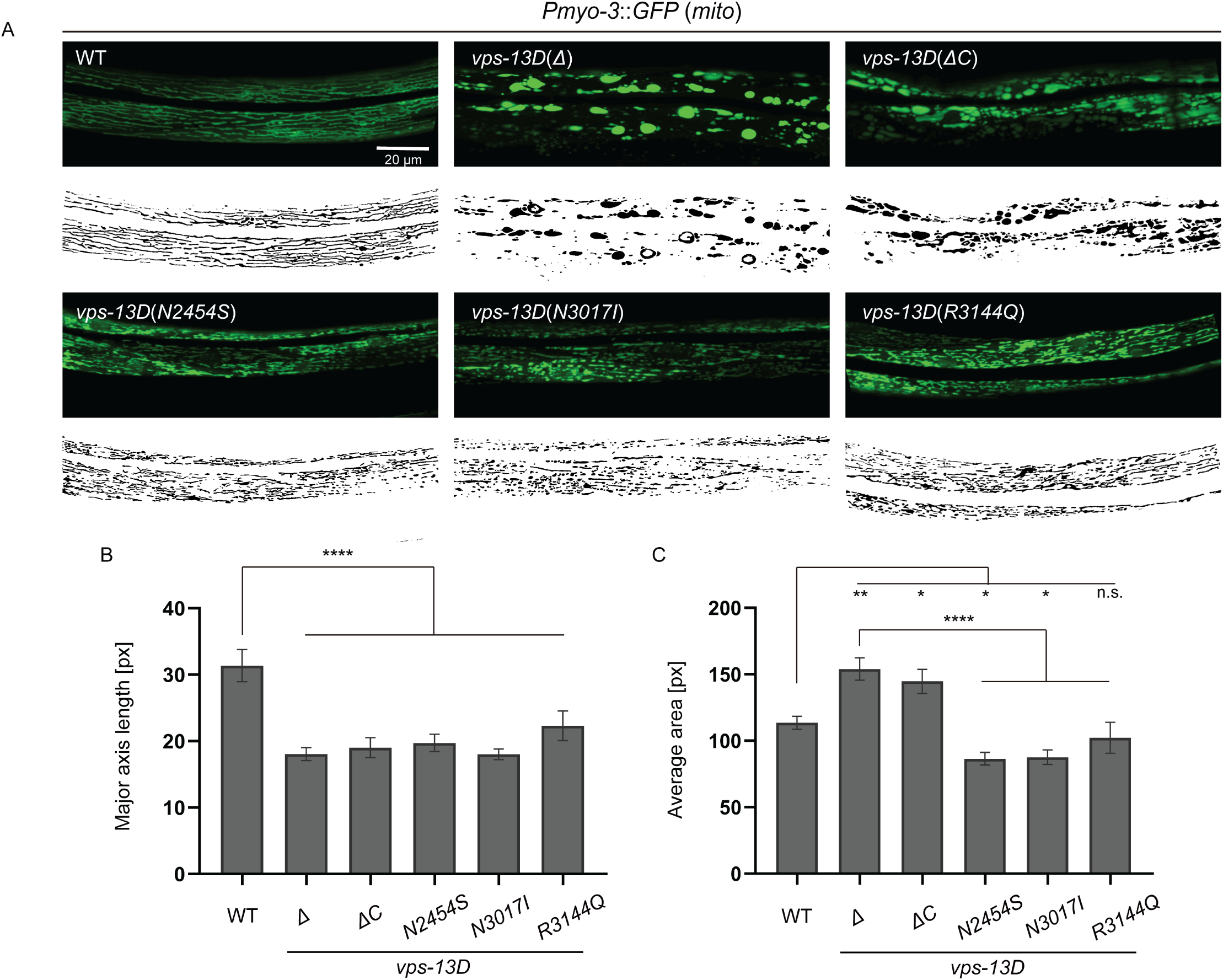
*vps-13D* regulates mitochondrial morphology in *C. elegans* (A) Mitochondrial morphology of the wild type and *vps-13D* mutant animals. Original images are at the top and MitoSegNet model segmentations are at the bottom. The scale bar is 20 μm. (B, C) Statistical analysis of mitochondrial morphology parameters. Major axis length (see materials and methods) and average area were measured in segmented images of mitochondria from wild-type and *vps-13D* mutant animals (n=15-25). One-way ANOVA with Dunnett correction was used to compare the difference between wild-type and mutant animals. Data is presented as mean ± S.E.M. *p <0.05, **p <0.01, *** p <0.001, **** p <0.0001, n.s. = not significant.

By contrast, *vps-13D*(*N2454S*) and *vps-13D*(*N3017I*) mutant strains display slightly smaller mitochondrial areas of 86.5±20.9 pixels and 87.6±26.5 pixels, respectively, where *vps-13D*(*R3144Q*) mutant is similar to the wild type (102.3±49.4 pixels) (Fig. 3A and 3C). These results indicate that SCAR4 mutations in the *C. elegans VPS13D* ortholog *vps-13D* lead to alterations in mitochondrial morphology, supporting VPS-13D’s role in maintaining mitochondrial structure.

### *vps-13D* disruption induces mitochondrial unfolded protein response

Although previous assays have examined the effects of *VPS13D* mutations on mitochondrial morphology, their impact on mitochondrial homeostasis remains unclear (Anding et al. 2018). The mitochondrial unfolded protein response (UPR^mt^) is a crucial cellular pathway that protects mitochondria from multiple forms of cell stress (Shpilka and Haynes 2018). To test whether *vps-13D* mutants affect UPR^mt^, we used a transgene of the transcriptional reporter *Phsp-6::mtHSP70::GFP* to monitor UPR^mt^ in *C. elegans* (Yoneda et al. 2004; Yang et al. 2022). Interestingly, *vps-13D*(*Δ*) and *vps-13D*(*ΔC*) mutant strains exhibit a significant induction of the UPR^mt^ with high levels of *Phsp-6::mtHSP70::GFP* intensity compared to wild-type animals at L4 larval age (Fig. 4A and 4B). While the relative fluorescence intensity of *vps-13D*(*N2454S*) and *vps-13D*(*R3144Q*) missense mutants were not significantly different from wild type, homozygous *vps-13D*(*N3017I*) mutant worms exhibited a slight increase in UPR^mt^ reporter expression compared to control worms (Fig. 4A and 4B). Combined, these results indicate that VPS13D plays a crucial role in maintaining mitochondrial homeostasis. Since the large deletion *Δ*(*ok2632*) allele, and the C-terminal deletion allele *ΔC* (*zf197*) causes similar deficiencies in fertility, development, locomotion, as well as mitochondrial morphology, we cannot exclude that frameshift mutations near the C-terminus could compromise protein stability.

**Fig. 4.**
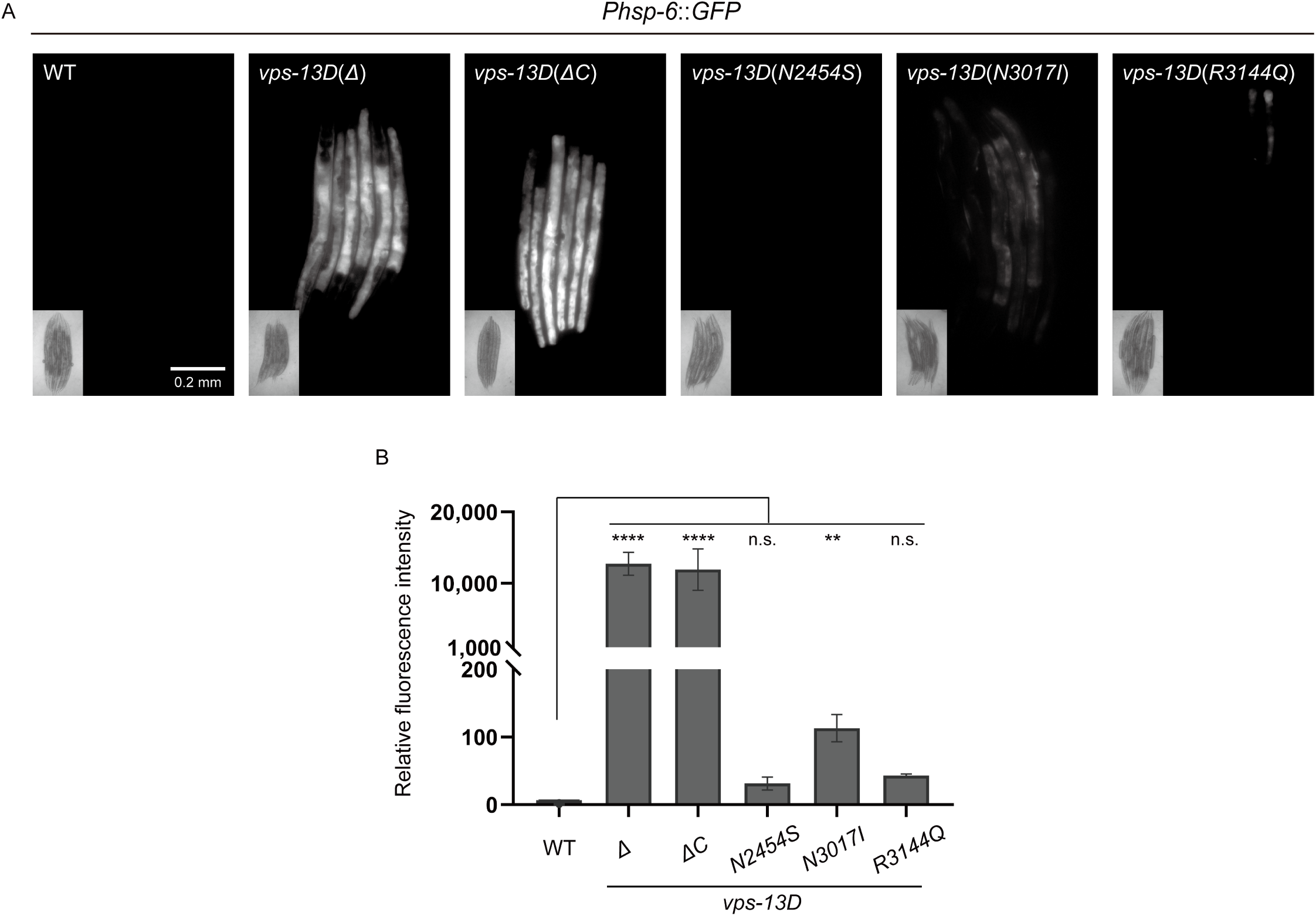
*vps-13D* dysfunction induces mitochondrial UPR (A) *Phsp-6*::*GFP* expression of wild type and *vps-13D* mutant worms. The scale bar is 0.2 mm. (B) Quantification of *Phsp-6*::*GFP* relative fluorescence intensity (n=3-5 groups*6 animals). Kruskal-Wallis H test with Dunnett correction was used to compare the difference between wild-type and mutant animals. Data is presented as mean ± S.E.M. *p <0.05, **p <0.01, *** p <0.001, **** p <0.0001, n.s. = not significant.

### *vps-13D* and *fzo-1/MFN2* function in a pathway to regulate mitochondrial homeostasis

In *Drosophila*, the orthologue of *MFN2*, *Marf* acts downstream of Vps13D to regulate mitochondrial fusion (Shen et al. 2021b). The *C. elegans* MFN2 orthologue FZO-1 is also involved in mitochondrial fusion (Rolland et al. 2009). Since our results indicate that *vps-13D* loss promotes alteration of mitochondrial morphology (Fig. 3), we examined the potential relationship between VPS-13D and FZO-1 in the regulation of mitochondrial morphology. We performed RNA-mediated interference (RNAi) targeting *fzo-1* either in *vps-13D*(*Δ*) mutant background or in control worms. *fzo-1* knockdown worms exhibited a decrease in average major axis length (17.7±2.5 pixels) and mitochondria area (77.8± 8.5 pixels) compared to control animals, which exhibited a major axis length of 28.4± 6.7 pixels and a mitochondrial area of 119.2 ± 9.8 pixels (Fig. 5A, 5B, and 5C). Knockdown of *fzo-1* failed to suppress the abnormal mitochondria morphology observed in *vps-13D*(*Δ*) deficient animals (Fig. 5A, 5B, and 5C). The major axis length (20.6±6.5 pixels) and mitochondria area (146.7±57.4 pixels) of *vps-13D*(*Δ*); *fzo-1*(RNAi) worms were similar to those observed in *vps-13D*(*Δ*) control animals (19.4±4.2 pixels in major axis length and 157.4±40.1 pixels in area). We also examined whether *fzo-1* affects mitochondria unfolded protein response (UPR^mt^). Consistent with previous studies (Haeussler et al. 2020), the knockdown of *fzo-1* in wild-type worms led to a significant increase in the level of the UPR^mt^ reporter *zcIs13* (Fig. 5D and 5E). However, *fzo-1* inactivation in *vps-13D*(*Δ*) mutant worms did not result in a significant change in UPR^mt^ compared to the *vps-13D*(*Δ*) single mutant strain (Fig. 5D and 5E). Collectively, these data suggest that *vps-13D* and *fzo-1* function together to regulate mitochondria homeostasis.

**Fig. 5.**
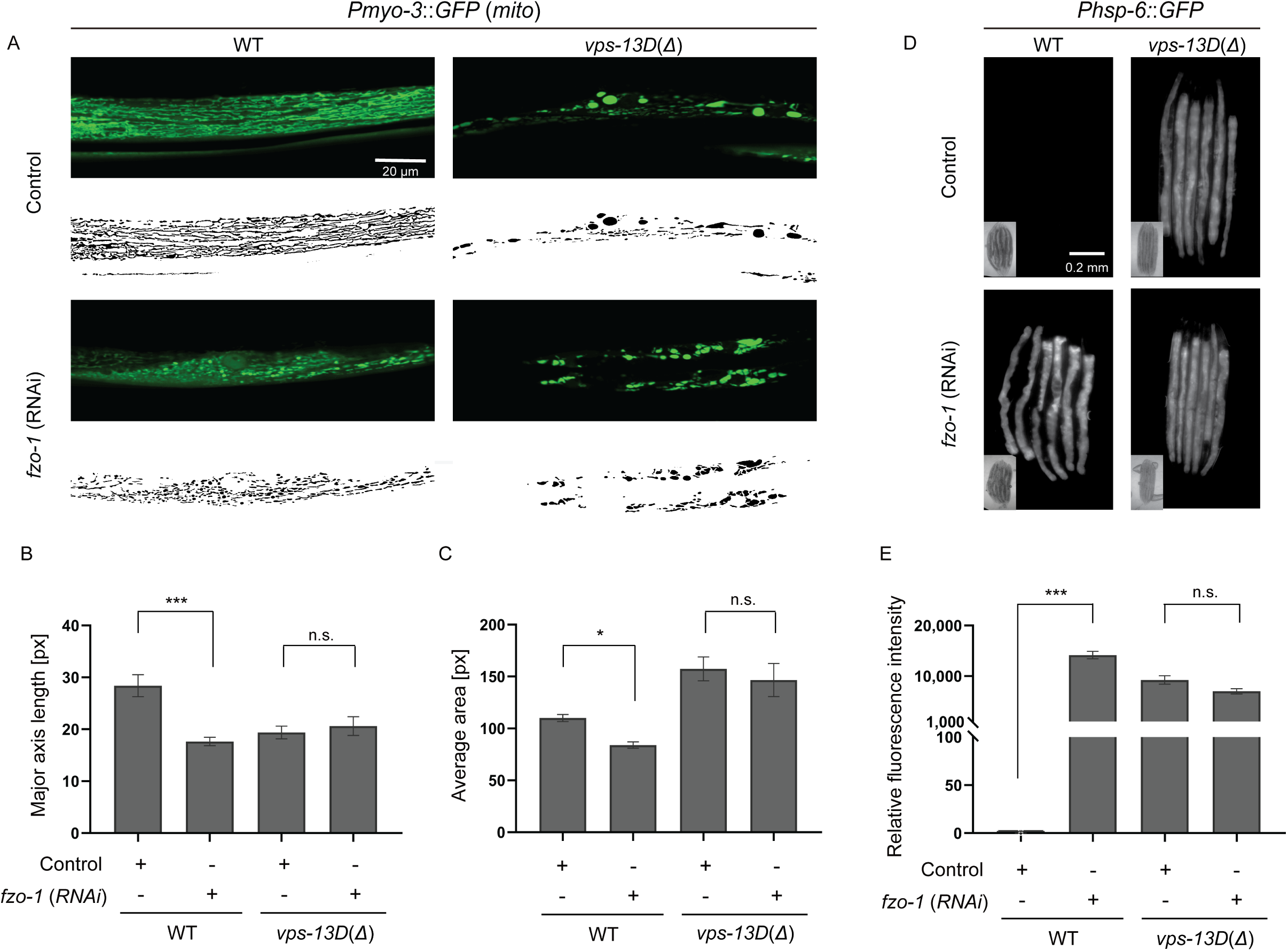
*vps-13D* and *fzo-1* regulate mitochondrial homeostasis in similar pathways (A) Mitochondrial morphology changes of wild type and *vps-13D*(*Δ*) animals grown on either control mock or *fzo-1* (*RNAi*). Original images are at the top and MitoSegNet model segmentations are at the bottom. The scale bar is 20 μm. (B, C) Statistical analysis of mitochondrial morphology parameters. Major axis length (see materials and methods) and average area were measured in segmented images of mitochondria from wild type and *vps-13D*(*Δ*) animals grown on either control mock or *fzo-1* (*RNAi*) (n=10-15). (D, E) *Phsp-6*::*GFP* expression of wild type and *vps-13D*(*Δ*) animals grown on either control mock or *fzo-1* (*RNAi*) (n=3-15 groups*6 animals). The scale bar is 0.2 mm. Kruskal-Wallis H test with Dunnett correction was used to compare the difference between wild-type and mutant animals. Data is presented as mean ± S.E.M. *p <0.05, **p <0.01, *** p <0.001, **** p <0.0001, n.s.= not significant.

## Discussion

We developed a new model to study *VPS13D* related SCAR4 disease in the nematode *C. elegans*. Our results indicate that *C. elegans* carrying the *VPS13D* deletions and disease-associated missense mutations, in the worm orthologue *vps-13D,* are viable. However, *vps-13D* null deletion mutations that remove the C-terminal display maternal effect sterility. In addition, *vps-13D* deletions and N3017I missense mutant *C. elegans* exhibit impaired locomotion, thus reflecting a similar role for this gene because human SCAR4 patients exhibit movement deficiencies. Importantly, *vps-13D* mutant *C. elegans* displayed abnormal mitochondrial morphology, and *vps-13D* deletions and *vps-13D*(*N3017I*) mutants displayed varying levels increased mitochondrial UPR.

The key clinical features of SCAR4 patients include progressive development of hyperkinetic movement disorder (dystonia, chorea, and/or ataxia). SCAR4 patient muscle biopsies exhibited mitochondrial accumulation and mild lipidosis (Gauthier et al. 2018). Similarly, altered mitochondrial morphology and mitochondria clearance is decreased in cultured *VPS13D* patient-derived fibroblasts, *VPS13D* mutant human HeLa cells, and *Drosophila* (Seong et al. 2018; Anding et al. 2018; Shen et al. 2021b). Although mitochondrial respiratory chain function was normal in the muscle biopsy of one SCAR4 patient, a decrease in energy production and complex (I, III, and IV) protein levels has been detected in some patient fibroblasts (Gauthier et al. 2018; Seong et al. 2018; Durand et al. 2022). Consistent with the movement defect of SCAR4 patients, worms carrying either deletions or N3017I missense mutations in *vps-13D* exhibited defects in crawling and/or swimming behavior. Interestingly, the abnormal swimming ability of *vps-13D* mutant worms is suggestive of mitochondrial dysfunction since swimming has been shown to be more energetically demanding than crawling for *C. elegans* (Laranjeiro et al. 2017; Kang et al. 2024). The similarities between SCAR4 patients and *vps-13D* mutant worms, including locomotion defects, mitochondrial morphology, and mitochondrial homeostasis changes, suggest that *C. elegans* will likely be a useful model to reveal other genes in this pathway because of the strength of this genetic model.

Mitochondrial homeostasis is controlled by mitophagy machinery, including the PINK1-Parkin axis (Pickrell and Youle 2015). VPS13D functions downstream of PINK1, but bifurcates with Parkin to regulate mitochondrial clearance (Shen et al. 2021a). Specifically, VPS13D binds to ubiquitin and affects both the PINK1 substrate ubiquitinphospho-serine 65 and localization of the mitophagy receptor Ref2p/p62 on mitochondria (Shen et al. 2021a). Previous studies suggest mitophagy and mitochondrial UPR are both required to maintain mitochondrial health (Pellegrino and Haynes 2015). Mitophagy and mitochondrial UPR function in a complementary manner to either recycle the damaged mitochondria by autophagy or recover the less damaged mitochondria. Additionally, mitophagy inhibition also causes damaged mitochondrial DNA accumulation that triggers mitochondrial UPR. Recently, it has been shown that non-mitochondrial proteins, including VPS13D, also play a role in maintaining mitochondrial homeostasis (Rolland and Conradt 2022). Although it is unclear if VPS13D regulates mitochondria clearance in worms, evidence of VPS13D regulating mitophagy in different animals and human cells suggest this possibility (Anding et al. 2018; Shen et al. 2021a). Therefore, *vps-13D* mutant worms may potentially accumulate damaged mitochondria to activate mitochondrial UPR.

Mitochondria and ER contact dictates inter-organelle lipid transfer sites and mitochondrial membrane lipid composition (Gatta and Levine 2017). The lipid composition of mitochondrial membranes, in turn, can influence crucial processes within mitochondria (Bockler and Westermann 2014; Lahiri et al. 2015). Previous studies indicate Vmp1 and Vps13D regulate mitochondria-ER contact and mitophagy (Shen et al. 2021b). VPS13D binds the outer mitochondrial membrane GTPase-Miro (both Miro1 and Miro2), likely via the WD40-like/VAB domain) (Guillen-Samander et al. 2021), and the conserved C-terminal regions have been shown to work synergistically with the VAB domain to facilitate the membrane targeting (Dziurdzik and Conibear 2021). Moreover, the VPS13D C-terminus is similar to the lipid transfer protein ATG2 (Oztop-Cakmak et al. 2022). Consistent with the importance of these domains, our data indicate that *vps-13D* C-terminus is important for mitochondrial homeostasis.

The mitochondrial protein mitofusin-2 (MFN2) facilitates mitochondrial fusion, as well as mitochondria and ER contact (Filadi et al. 2018). Mutations in *MFN2* are associated with Charcot-Marie-Tooth disease type 2A (CMT2A), in which mitochondrial morphology is abnormal in nerve tissue and fibroblasts from patients (Verhoeven et al. 2006; Amiott et al. 2008). Similarly, loss of *fzo-1*, the worm *MFN2* homolog, resulted in progressive movement deficits, fragmented mitochondria, and mitochondrial UPR activation in *C.elegans* (Byrne et al. 2019; Haeussler et al. 2021). Consistent with these studies, our results indicate *fzo-1* knockdown worms displayed altered mitochondrial morphology and induced mitochondrial UPR. UPR^mt^ activation requires the key transcription factor ATFS-1, which regulates the *hsp-6* reporter expression (Lin et al. 2016; Xin et al. 2022; Shpilka et al. 2021; Nargund et al. 2015). Notably, knockdown of *atfs-1* significantly suppressed UPR^mt^ in the *fzo-1* mutant worms, confirming that UPR^mt^ induction by loss of *fzo-1* is ATFS-1 dependent (Chen et al. 2021). Unlike *Marf/MFN2* function in the Vps13D pathway regulating mitochondrial morphology in *Drosophila* (Shen et al. 2021b), our results suggest that *fzo-1* and *vps-13D* could function in a related pathway to affect mitochondrial UPR in *C. elegans*. In summary, this study provides evidence supporting the conserved role of VPS13D in regulating mitochondrial morphology and function in *C. elegans* and provides insights how disease mutations affect mitochondrial homeostasis and behavior.

## Acknowledgments

We thank the Caenorhabditis Genetics Center (CGC), which is funded by the NIH Office of Research Infrastructure Programs (P40 OD010440), for some worm strains. We thank W.K. Kang, J.T. Florman, and M. Gorczyca for technical support and helpful discussions.

## Funding

This work was supported by National Institutes of Health grants R01 GM140480 (MJA) and R35 GM131689 (EHB).

## Conflicts of interest

The authors declare no conflict of interest.

## Data availability

All data files from this study have been uploaded to Mendeley Data (DOI: 10.17632/3mwgvd2dbx.1).

## Figure legends

**Fig. S1.** Day 1 *vps-13D* mutant worms did not show a significant deficiency in locomotion.

(A) Comparison of average velocity for 1-day-old young adult wild type and *vps-13D* mutant animals from the final 5 mins MWT recordings. The velocity has been normalized and calculated based on the worm body lengths of respective strains. The difference in average velocity between wild-type and mutants was analyzed by one-way ANOVA with Dunnett’s multiple comparison test.

(B) Number of thrashes per minute in M9 liquid for 1-day-old young adult wild type and *vps-13D* mutant animals (n=10-15). Differences between wild type and mutants were analyzed by One-way ANOVA with Dunnett correction for multiple comparisons.

